# Kombucha as a model system for multispecies microbial cooperation: theoretical promise, methodological challenges and new solutions ‘in solution’

**DOI:** 10.1101/214478

**Authors:** Alexander Niall May, James Medina, Joe Alcock, Carlo Maley, Athena Aktipis

**Author notes:** At the time of this writing, James Medina is at Washington University, St. Louis, MO. Address correspondence to: Alexander N. May (, 1-480-930-7796, ORCID ID: 0000-0003-4521-6916) or Athena Aktipis.

## Abstract

Kombucha is a sweetened tea fermented by bacteria and yeast into a carbonated, acidic drink, producing a surface biofilm pellicle (colloquially called a SCOBY) during the process. Typically, liquid and a biofilm pellicle from a previously fermented culture is used as a starter for new cultures; however, there is no standard protocol for growing kombucha in the laboratory. In order to establish a standard protocol with low variability between replicates, we tested whether we could begin a kombucha culture with only well-mixed liquid stock. We found that viable kombucha cultures can be grown from low percentages of initial inoculum stock liquid, that new pellicles can form from liquid alone (with no ‘starter’ pellicle), and that the variation in the pellicle characteristics is lower when only a liquid starter is used (p = 0.0004). We also found that blending the pellicle before including it significantly reduces the variation among replicates, though the final pellicle was abnormal. We conclude that growing kombucha from only liquid stock is viable and provides a greater degree of experimental control and reproducibility compared to alternatives. Standardizing methodologies for studying kombucha in the lab can facilitate the use of this system for exploring questions about the evolutionary, ecological and cooperative/competitive dynamics within this multi-species system including resource transfers, functional dependence, genetic divergence, collective defense, and ecological succession. A better understanding of kombucha and other fermented foods may eventually allow us to leverage their pathogen inhibitory properties to develop novel antibiotics and bacteriocins.

## Introduction

Kombucha is a sweetened tea that is fermented by bacteria and yeast into a carbonated, acidic drink, and it produces a cellulose biofilm pellicle (colloquially known as a SCOBY, or Symbiotic Community Of Bacteria and Yeast) on the surface of the kombucha liquid. It is purported to have originated in the Tsin Dynasty of Ancient China and then spread to Asia, Eastern Europe and Russia at the turn of the last century [1]. Kombucha is known by many names, including tea fungus, *Cainiigrib, Cainii kvass, Japonskigrib, Kambucha, Jsakvasska, Heldenpilz, Kombuchaschwamm,* and *Funkochinese* [2]. Kombucha is an excellent candidate model system for studying multi-species cooperation because it has a long history of being artificially selected by humans, it is non-toxic, it is easy to grow and grows quickly. In addition, its popularity makes an effective tool for both teaching and for engaging citizen scientists.

Kombucha has several properties that make it useful as a model system for social interactions in microbial ecosystems: its microbial composition has been characterized, the kinetics of its fermentation have been established, it has innate antimicrobial properties, and its culture conditions are simple. However, there are currently several methodological challenges to using kombucha as a model system, including the need to control initial conditions. Here we describe the promise of kombucha as a model system, the nature of those methodological challenges, and describe results of experiments showing that starting replicates ‘in solution’ without a pellicle produces more consistent resulting pellicle sizes compared to existing protocols that include a piece of pellicle in the starting conditions. We also tested the effects of blending the pellicle in the inoculum.

Multispecies ecological systems abound in the natural environment, and are also an important part of many human societies and cultures in the form of fermented foods and beverages. The microbial communities in these fermented foods are typically well-characterized, easy to maintain, and highly reproducible; they represent a bridge between fully-artificial *in vitro* systems and far more complex natural ecosystems.

Natural multispecies communities are enormously rich in diversity, but untangling the multitudes of relationships and interactions is complex, even with modern molecular tools. Additionally, many wild species are difficult or impossible to isolate or culture, restricting the ability to manipulate and test hypotheses about their functional hierarchy *in vitro* [3]. On the other hand, artificial microbial systems in the laboratory provide powerful microcosms to test fundamental ecological concepts within tightly controlled parameters, usually using well-characterized species. The downside of artificial microbial model systems is that they may not be as applicable to natural microbial systems. Artificial microbial models systems often have other limitations including restricted scalability, unique microbial life histories that may or may not be shared with natural systems, and a rapid speed of evolutionary change that can introduce challenges for characterizing the dynamics among microbes within these systems [4].

Kombucha has many of the advantages of natural microbial systems without the disadvantages of many artificial laboratory systems. Kombucha is a natural (albeit artificially selected) multispecies microbial system that can be easily cultured in controlled laboratory conditions. With its unique bacterial-fungal symbiosis, biofilm-forming properties, and simple propagation, Kombucha may provide opportunities for answering questions that cannot be answered with artificial microbial systems or with completely wild microbial systems. Fermented foods in general have been domesticated by humans to grow in culture conditions, originally for human consumption. Many of the characteristics that make fermented foods easy to cultivate in homes and kitchens also make them excellent candidates for growing in the laboratory. For these and other reasons, fermented foods are now beginning to be used more broadly as alternative model systems for microbial ecosystems [3].

Typically, kombucha is cultured over a 7–14 day period at room temperature. Acetic acid bacteria in this community produce a thick cellulose biofilm pellicle (colloquially known as a SCOBY, or Symbiotic Community Of Bacteria and Yeast) on the surface of the solution, in which bacteria and yeast are embedded [5]. Fermentation begins with invertase production by yeast, which may act as a public good by cleaving sucrose into glucose and fructose (both of which are utilized by both yeast and bacteria, Fig 1). Then the bacteria produce an encapsulating surface biofilm via cellulose polymerization of glucose monomers. Each successive round of fermentation typically adds a new layer of pellicle to the surface of the solution, creating a multi-layered pellicle over time (Fig 2). This biofilm may also act as a public good through the creation of a barrier to microbial invasion, among other putative benefits, such as: retaining buoyancy, providing a substrate for colonization, competitor exclusion, water retention, and resistance to UV rays [6]. The bacterial-yeast symbiosis appears to be characterized by a reciprocal benefit with yeast liberating resources and bacteria protecting the system from invasive microbial species. Despite kombucha’s inherent ease of replication and culturing, there is no existing standard protocol for generating new cultures from pre-existing stocks or collected samples. Typically, liquid and a biofilm pellicle from a previously fermented culture is used as a starter for new cultures; however, there is often significant variation in the amount of liquid and biofilm used in these studies [2; 7]. Additionally, the biofilm itself is a highly spatially-structured community with uncertain consistency and unknown population densities. Therefore the use of a biofilm pellicle in starting new cultures may introduce unnecessary variation into the stock, resulting in low experimental control and low reproducibility.

**Figure 1.**
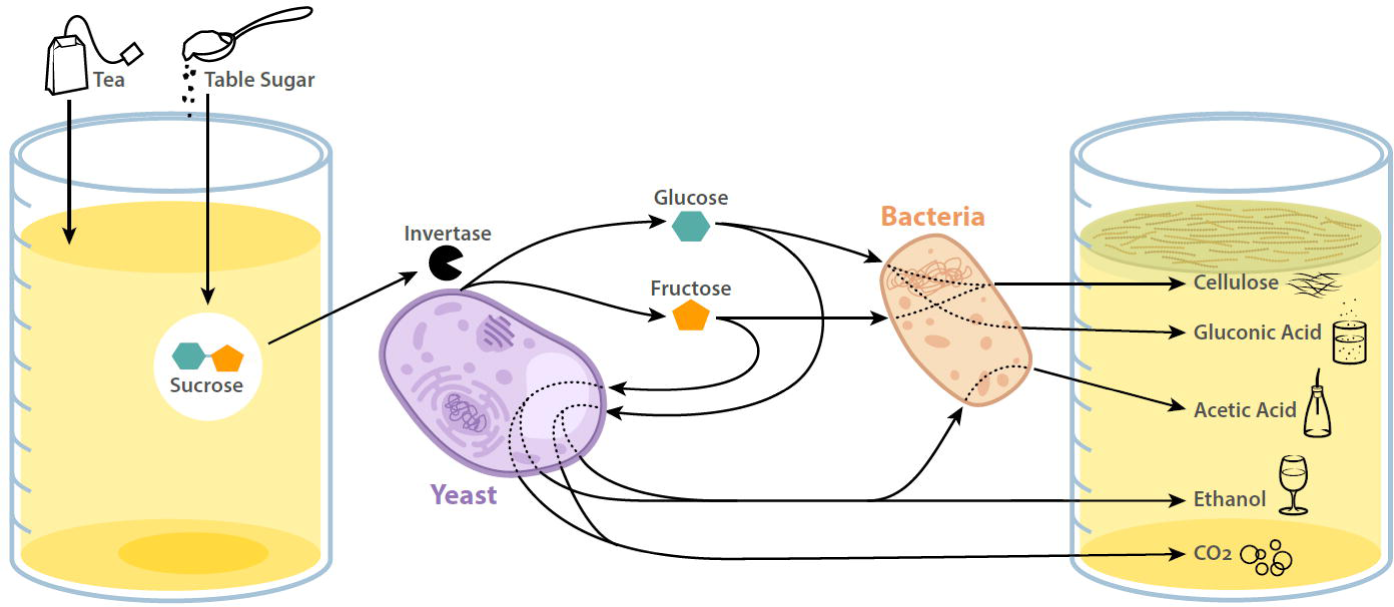
(made with Adobe Illustrator). Kombucha is the result of a symbiotic association between acetic acid bacteria and ethanol-fermenting yeast, where the yeast cleave sucrose into glucose and fructose via invertase in the cell wall, which then acts as public goods for both yeast and bacteria. Bacteria produce acid, ethanol, carbon dioxide as well as a cellulose biofilm that forms a barrier on the top of the solution.

**Figure 2.**
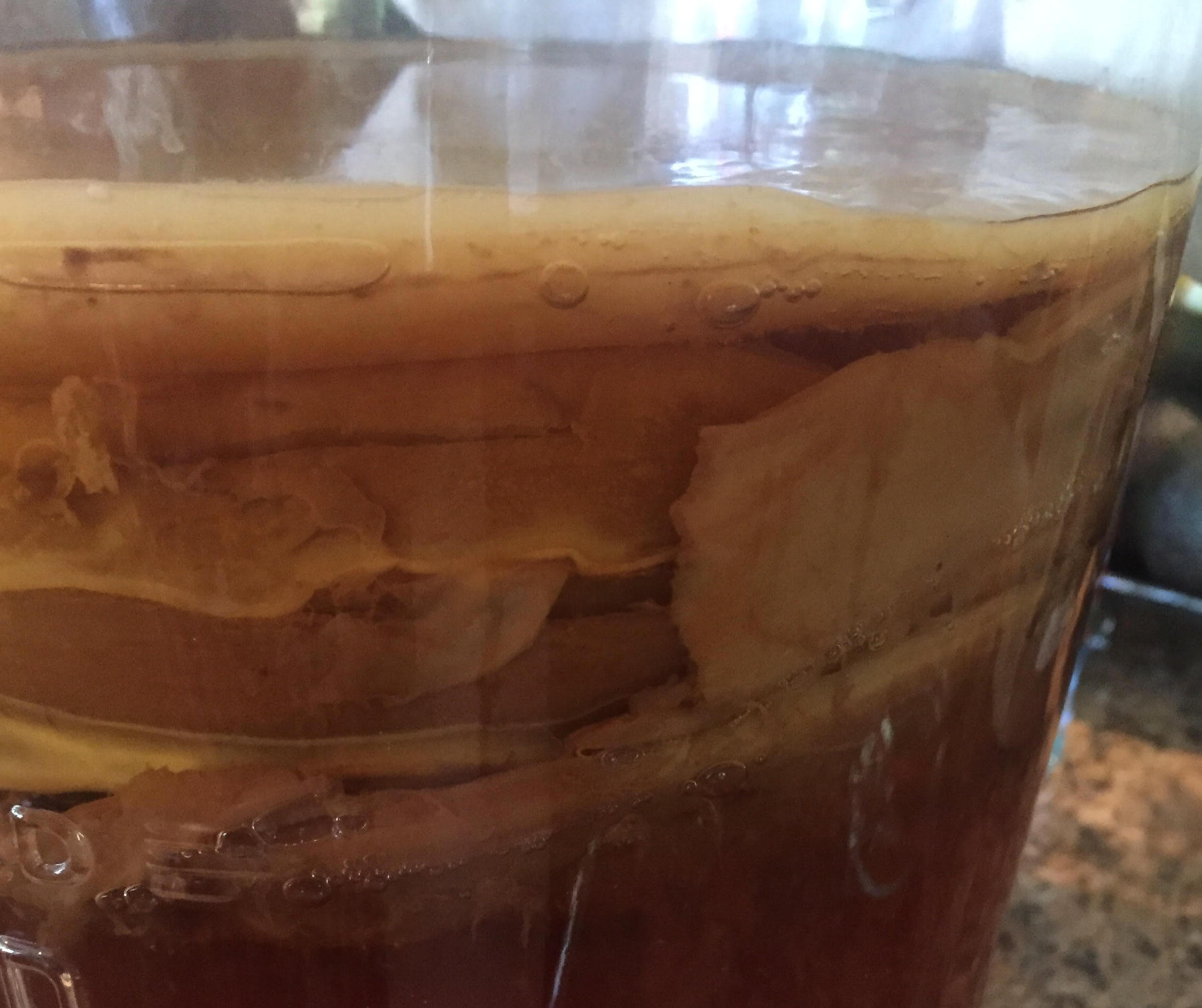
(photograph). A typical kombucha ‘stock’ solution as it appears during growth at room temperature. If allowed to ferment continuously, new pellicle layers will form at the liquid-air interface upon addition of new media, forcing the older layers to submerge beneath. Strand-like material is often seen dangling from the upper levels of pellicle into the liquid phase.

### Results

To address these reproducibility issues and develop standard protocols for growing kombucha, we ran several experiments to determine whether a well-mixed liquid inoculum from a kombucha culture can be used alone (without an initial pellicle) to recreate the kombucha culture. We examined this under three levels of initial liquid inoculum amounts (5%, 10% and 15%), testing whether this initial inoculum mixed with tea could give rise to a new pellicle (from only the solution without initial pellicle) and also whether cultures that are started with only the inoculum solution had lower variance in the weight of the pellicle that formed on the surface after 18 days, compared to those cultures that were started with both inoculum and an initial pellicle.

We discovered that, contrary to documented practice, it is possible to start kombucha from stock alone, without any pellicle. The size of the resulting pellicle depends on the proportion of stock in the inoculum (Experiment 1; Fig 3 and Table S1). There was no statistically significant difference in the resulting pellicle weight when comparing kombucha started from stock alone versus stock with a pellicle fragment (T-Test p>0.05 for all comparisons; Table S2). However, using a fragment of pellicle in the starter led to much higher variance in the resulting pellicles, compared to stock alone (Levene’s test for unequal variances *p*=0.0004; 1-way ANOVA: F(1,16)=21.85, *p*<0.00025).

**Figure 3.**
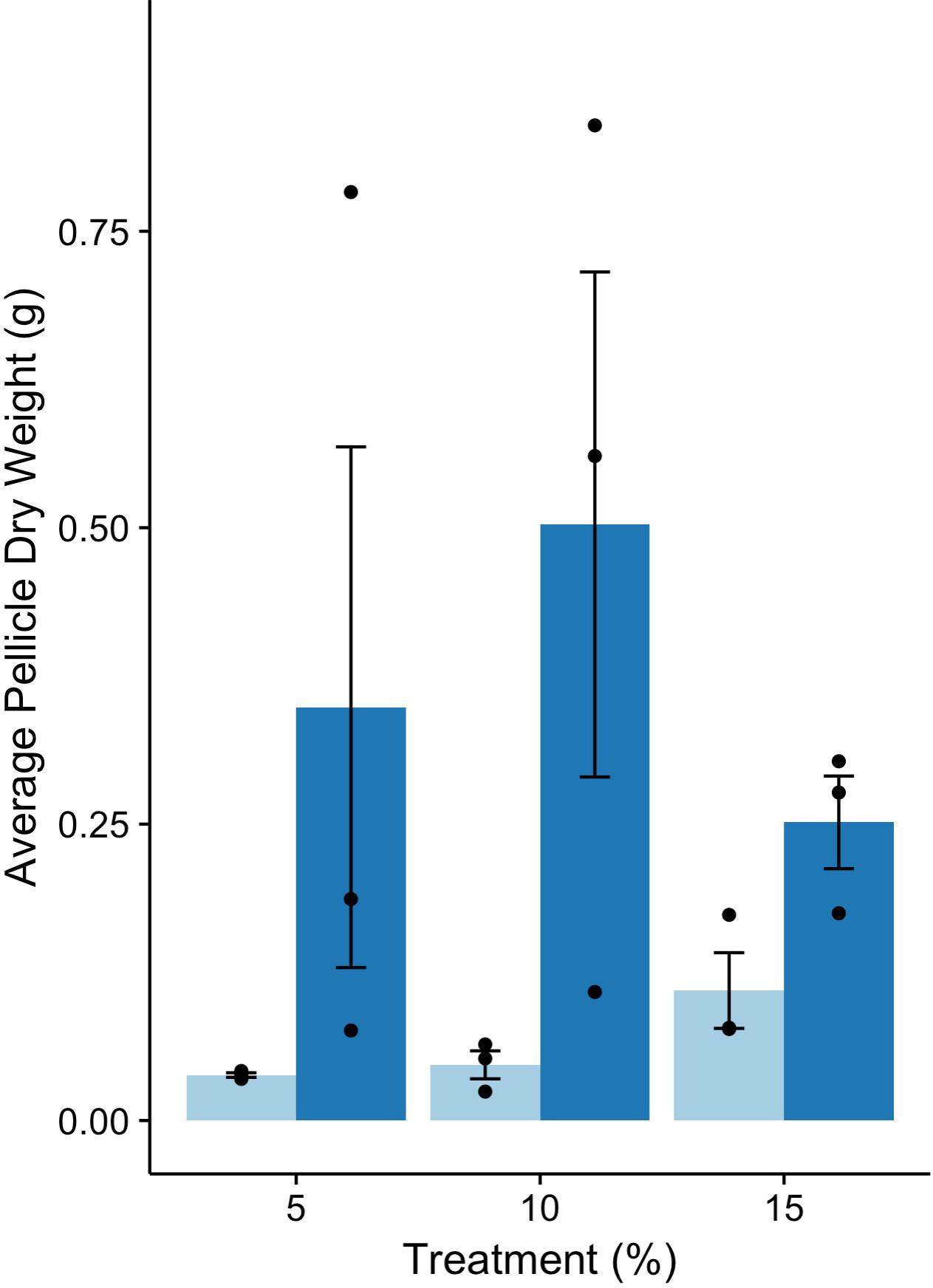
(made with R software). Average of dried pellicle weights across each treatment (treatments with light blue bars contain only liquid, while treatments with dark blue bars contain liquid plus pellicle fragments; the thin capped black bars represent standard error of each average and individual replicate data points are represented as black dots over the treatment bars). The treatments are: 5% starting stock with sweetened tea (5% Stock), 5% starting stock with sweetened tea and 3g pellicle fragment (5% Stock + Pellicle), 10% starting stock with sweetened tea (10% Stock), 10% starting stock with sweetened tea and 3g pellicle fragment (10% Stock + Pellicle), 15% starting stock with sweetened tea (15% Stock), and 15% starting stock with sweetened tea and 3g pellicle fragment (15% Stock + Pellicle).

Next, we collected 0.5 grams of pellicle per treatment by randomly sampling locations in a pellicle with a 10mm punch biopsy. This collection of biopsies was either blended (homogenized) or left as a collection of intact pellicle fragments (fragmented) and then added to the inoculum. We predicted that starting with a blended (homogenized) pellicle would lead to lower variance in the ending dry weights because the spatial variation in the pellicle structure would be eliminated (compared to fragments which would maintain some variability at the microstructure level). As predicted, we found that the blended treatment had low variance compared to the fragmented condition (Fig 4 and Table 1) and that the variance in the ending weights of the blended and liquid-only condition were equal (Levene’s test p=0.06583). When analyzed overall, the groups had unequal variances (Levene’s test *p*=5.284x10**^-05^**), and that all other pairwise comparisons (other than the liquid-only vs. blended treatment) had unequal variance (Table 1; column 1). Natural-log transformation of these data did not abrogate the variance inequality between treatments (Table 1; column 2).

**Figure 4.**
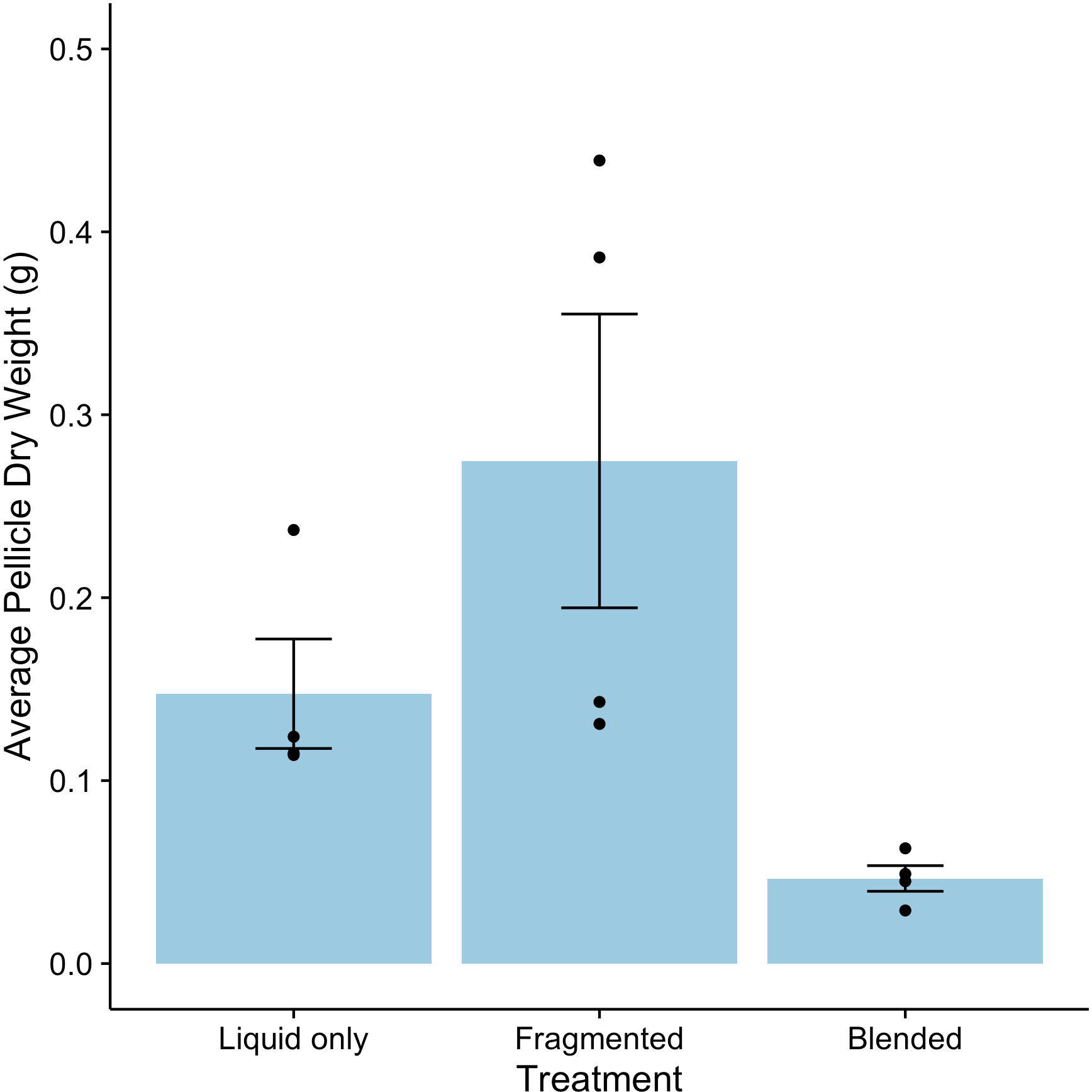
(made with R software). Average of dried pellicle weights comparing physical treatment of the pellicle fragment in the inoculum (the large light blue bars are the averages of each treatment, the thin capped black bars represent standard error of those averages, and individual replicate data points are represented as black dots over the treatment bars). The treatments are 10% starting stock with sweetened tea (LiquidOnly), 10% starting stock with sweetened tea and 0.5g of pellicle fragments (Fragmented), and 10% starting stock with sweetened tea and 0.5g blended pellicle fragments (Blended).

We found that either blending or fragmenting the starter pellicle fragment had a significant effect on the resulting dry pellicle weight in experiment 2 (1-way ANOVA: F(2,9)=13.68, *p*=0.00187, Fig 4). Blending the starter pellicle fragments led to a lighter pellicle than starting from stock alone (post-hoc Tukey test, p=0.0166), and when starting from intact pellicle fragments (post-hoc Tukey p=0.0014). There was no significant difference in pellicle weight between conditions that were started with stock alone versus stock and intact pellicle fragments (post-hoc Tukey p=0.2875), in agreement with experiment 1. The blended pellicle treatments produced lighter pellicles with a translucent appearance that quickly fell apart when handled. The structure of these blender-derived pellicles was gelatinous and runny, unlike the firm pellicles produced by the liquid-only and liquid-plus-fragments treatments. This abnormal appearance and structure is a concern for using this blended protocol for future work (see discussion) despite the fact that blending led to lower variance among replicates than other methods involving including the pellicle.

## Discussion

The inherent diversity of species, sources, and conditions used to cultivate kombucha pose challenges for its development as a model system for multispecies interactions. We have shown that an ‘in solution’ approach using only liquid inoculum produces kombucha pellicles with significantly less variance in the dry weights than those produced with liquid and a supplemental pellicle inoculum. This suggests that the use of a pellicle in current kombucha culturing practices may introduce unnecessary variation in outcomes between conditions. Our results also suggest that the physical structure of the pellicle may contribute to variance among replicates: when we used a blended pellicle (SCOBY), this resulted in less variance among replicates. When we used large fragments of the pellicle, we found high variance among replicates, suggesting that the microstructure (smaller than the 0.5 gram fragments) may contribute to the variation among the replicates.

We also showed that the variance in final pellicle weight increases with the percent of stock in the inoculum. Based on these results, we suggest using a 10% stock with no pellicle for the inocula for follow-up studies because this amount of stock offers a reasonable growth rate without excessive variance.

### Historical culturing of kombucha and problems of origin

Owing to its unstandardized historical transmission, the details of how to propagate kombucha cultures are mostly anecdotal; there is no universal methodology for growth. Nevertheless, the traditional practice is to use a whole pellicle and a portion of liquid from a previously fermented batch, placing both into freshly brewed tea supplemented with sucrose (table sugar). The amount and type of each ingredient and the precise timing of brewing and fermentation vary with locality [2], which may contribute to the diverse microbiological communities found therein [5; 8; 9]. While this diversity is intriguing from an ecological standpoint, it represents a problem for reproducibility: every potential origin of kombucha is composed of different founding genera, species, and strains, and has evolved under different environmental selection pressures. These characteristics of kombucha create challenges for generalizing from results from specific studies of kombucha. Some of these challenges can be addressed by standardizing growth conditions and the initial stock.

### Spatial structuring and diversity of the community

Kombucha has additional complexities arising from the fact that there are at least 2 spatially distinct regions within each culture: the pellicle and the liquid (and possibly even a third distinct region: the interface between the pellicle and liquid, which has tentacle-like strands descending from the pellicle to several inches in the liquid, Fig. 2). These regions are known to differ in composition of their community members [5; 9] - an unsurprising result, considering the microenvironmental structural differences.

Intriguingly, Teoh et al. [10] found that the populations of certain yeast species declined in the liquid solution after 8-10 days of fermentation (as pH fell and nutrients were depleted), yet were able to persist at stable levels within the pellicle beyond this timeframe. The biofilm matrix may thus act as an encapsulating barrier, protecting the entire community against the increasingly harsh environmental changes, and allow otherwise intolerant species to remain as members of the system. This may be one of the constraints of using a liquid-only starter: by starting from kombucha liquid alone, we may miss certain species that can survive in the biofilm matrix but not the liquid.

Current kombucha culturing practice utilizes both liquid and pellicle regions as inoculum for new kombucha cultures. Our results suggest that this method may contribute to unpredictability of growth. The amount of the pellicle and its age likely contribute to the starting population size and diversity, both of which will have an effect on future development and community structuring. Unless the amount of pellicle inocula is carefully controlled and the composition is well-characterized, it is likely that the heterogeneity present in the microstructure will bias the development of the culture. In our study, including pellicle fragments did indeed generate larger variance in pellicle dry weight than was the case with their stock-only treatment counterparts (Table 1).

Previous work suggests that the liquid medium may have lower yeast population number per unit volume [10]. Despite this, we found that using well-mixed kombucha liquid without pellicle resulted in less variance in dry weight. Reports are inconsistent regarding whether the liquid has less diversity in yeast composition than the pellicle [5] or more diversity in yeast composition than the pellicle [9], which makes it difficult to contextualize our results in terms of the potential influence of yeast diversity on the outcomes we observed.

### Addition of pellicle as inoculum can add variability to future pellicles

Despite these advantages of using well-mixed liquid without pellicle, it is important to note that liquid-only culture did result in a lower total weight of the pellicle compared to liquid-and-pellicle cultures. This means that, if the goal is producing a large pellicle, starting with an initial pellicle may be a good practice. Our results do not suggest that use of initial pellicles is misguided in general, only that they may add variability in laboratory conditions that could be better controlled by starting only with well-mixed initial stock. Further, our results do not suggest that culturing practices of kombucha in the home or in commercial situations should be changed so as to not include an initial pellicle. Though they do suggest that including an initial pellicle may introduce more variability in the outcome of the resulting pellicle that grows on the surface of the liquid.

### Impact of blended homogenization of pellicle inoculum on new cultures

One of our conditions was to fully homogenize a pellicle in a blender and add 0.5 grams of the resulting material. We were curious as to whether including homogenized pellicle could reduce the variance and perhaps provide an alternative to liquid-only conditions if experimenters wanted to include pellicle in the initial conditions. Variance was indeed lower when a blended pellicle was used as inoculum as opposed to a fragment of pellicle. In fact, the variance when using a blended pellicle was no different than the liquid-only condition, suggesting that a blended pellicle may be a reasonable alternative to liquid. However, we also discovered that this fully blended pellicle produces a new pellicle that lacks structural integrity and has reduced dry weight compared to a non-blended fragment or a liquid-only condition. Whereas a typical pellicle resembles fruit leather in texture and consistency, the pellicle that resulted from the blended pellicle has a mucousy and flimsy structure that comes apart easily. Future work could help to distinguish between several possible explanations for this phenomenon. One possibility is that, when using a normal pellicle, the cellulose fibers in the pellicle act as a nucleus for the growth and organization of new fibers, like the seeding of a crystal. It may be the case that disrupting that organization by blending may generate many micro-seeds that do not connect well with each other as they grow, perhaps as a result of uncoordinated timing of the growth of bacterial cellulose and yeast hyphae that may contribute to the structural integrity. It is also possible that the blending protocol could have damaged the organisms responsible for biofilm formation, or caused the release of toxins into the liquid via cell lysis.

### Future directions for kombucha as a model system

While our experiments demonstrate repeatability in pellicle formation, much remains unanswered about culturing this system. Some researchers have investigated alternative carbon sources and substrates for the cultivation of kombucha (summarized by [2]), but little work has been done on the long-term adaptations that may occur from using non-canonical sugars and nitrogen sources. From an evolutionary perspective, this may provide clues as to the functional roles of the ecological partners in carbon acquisition, and could reveal unexpected evolutionary trajectories for future studies. As there is no universally recognized species composition for the fermentation process, it would be interesting to see whether convergent evolution has occurred across cultures due to the similarities in growth conditions. Likewise, the ability of kombucha to respond to environmental fluctuations could represent a powerful proxy for studying the behavior of natural systems. The introduction of antibiotics, antifungals, enzymes, inhibitors, neutralizing agents, toxins, or pollutants may all have unforeseen effects on the system that could be extrapolated to larger scale networks.

In addition to basic scientific inquiries, kombucha holds promise for more applied research. Its ability to inhibit pathogens has been studied i*n vitro* and is largely (though not exclusively) attributed to its acidic character. Greenwalt et al. [11] saturated cellulosic discs with solutions of fermented and unfermented black and green teas and found that kombucha cultures containing 7g/L of acetic acid (33g/L total acid) had an inhibitory benefit against an array of common human pathogens, though not against *Candida albicans*. Upon neutralization to a pH of 7, these benefits were abrogated even with increasing tea concentrations. In contrast, Sreeramulu et al. [12] demonstrated that a kombucha black tea solution retained inhibitory activity against *Escherichia coli, Shigella sonnei, Salmonella typhimurium, Salmonella enteritidus,* and *Campylobacter jejuni* even after neutralization to a pH of 7 or thermal denaturation at 80°C for 30 minutes. The exact nature of these inhibitory properties are unknown, but could include novel antibiotics or bacteriocins adapted to function under various environment conditions.

Unlike rationally designed therapeutics that can be thwarted by the evolution of resistance, the kombucha community itself can evolve and generate diverse molecules to kill competitors. Challenges to the system using invasive, human-associated pathogens could provide a platform for discovering new anti-microbial compounds. Additionally, the kombucha microbial community itself, while untested in human health, could provide an easy way to generate diversity within the gut microbiome. Tolerance to low pH would enhance survival of the microbes through the digestive tract, while the biofilm itself could be used as a source of nutrients for cellulose-degrading organisms during transit.

### Concluding remarks

There is a need for a thorough characterization of kombucha genetic and phenotypic diversity. The most comprehensive metagenomics study of kombucha to date included only 5 biofilm samples [5], and no studies have searched for viruses, despite viruses being the most abundant entity on the planet (10^31^ viral particles at any given moment) [13]. Further, resilience of the kombucha symbiotic community has not been systematically investigated: the minimal starting conditions, the ranges of growing conditions and the multispecies interactions required to prevent collapse are all currently unknown. Our experiments are a first step towards standardizing methodologies for kombucha that can facilitate the use of this system for exploring more evolutionary questions, such as the nature of resource exchange, functional dependence, genetic divergence, collective defense, and ecological succession.

## Methods

### Kombucha Media Preparation

In order to produce the sweetened tea media (STM), a recipe was developed based on the following protocol: 1L of deionized water (LabChem, PA) was brought to a boil (approximately 90°C) in a 1.7L stainless steel kettle (Hamilton Beach, NC), and then poured into a 1L Pyrex beaker containing 5g (0.5% w/v) of loose leaf black tea (Lipton, Unilever, UK) with 50g (5% w/v) of sucrose (Biobasic, NY). The solution was stirred until the sucrose was fully dissolved, and then allowed to steep for 10 minutes while the beaker was covered with aluminum foil. At this point, the solution was strained into a new Pyrex beaker through BrewRite coffee paper filter (Rockline Industries, WI) to remove tea leaf debris. To ensure uniformity across all replicates, all strained tea was pooled into a single graduated pitcher prior to distribution to replicate beakers. The tea was allowed to equilibrate to room temperature (21°C), but further sterilization steps were not performed in order to more accurately mimic the traditional processes used to prepare kombucha.

### Culturing Kombucha

To establish a laboratory-acclimated kombucha stock, 3 starter pellicles totaling approximately 1kg (fresh weight) were donated from Dr. Aktipis’s personal supply, along with approximately 500mL of kombucha liquid that had been propagated for an unknown length of time. This stock supply was placed into a 4 gallon (7.5L) glass jar (Nantucket Glassware, MA) along with 3.5L of freshly-prepared STM, bringing the total volume to 4L of liquid, plus 1kg of pellicle material. Two layers of approximately 150um-diameter pore cheesecloth (Carolina Biological Supply Company, NC) were placed over the top of the jar and secured with a rubber band. This stock solution was allowed to ferment continuously at room temperature, with removal and replenishment with approximately 1L of STM every four to six weeks. Additionally, whenever liquid was removed for experiments, an equal amount of STM was replenished into the stock solution. At the time of the experiment, the stock solution had been fermenting for 6 months at room temperature.

### Stock-Percentage Experiment

The goal of this experiment was to determine whether using liquid inoculum at different proportions with STM and in combination with biofilm fragments could produce replicable results during the generation of new kombucha cultures (Fig 5). To produce replicate kombucha vessels, liquid from the Kombucha stock solution was gently agitated for 15 seconds, and then combined with STM in 600mL Pyrex beakers in various proportions based on the treatment. These treatments consisted of 3 replicate beakers of 5% stock mix (380mL STM, 20mL stock), 10% stock mix (360mL STM, 40mL stock), and 15% stock mix (340mL STM, 60mL stock).

**Figure 5.**
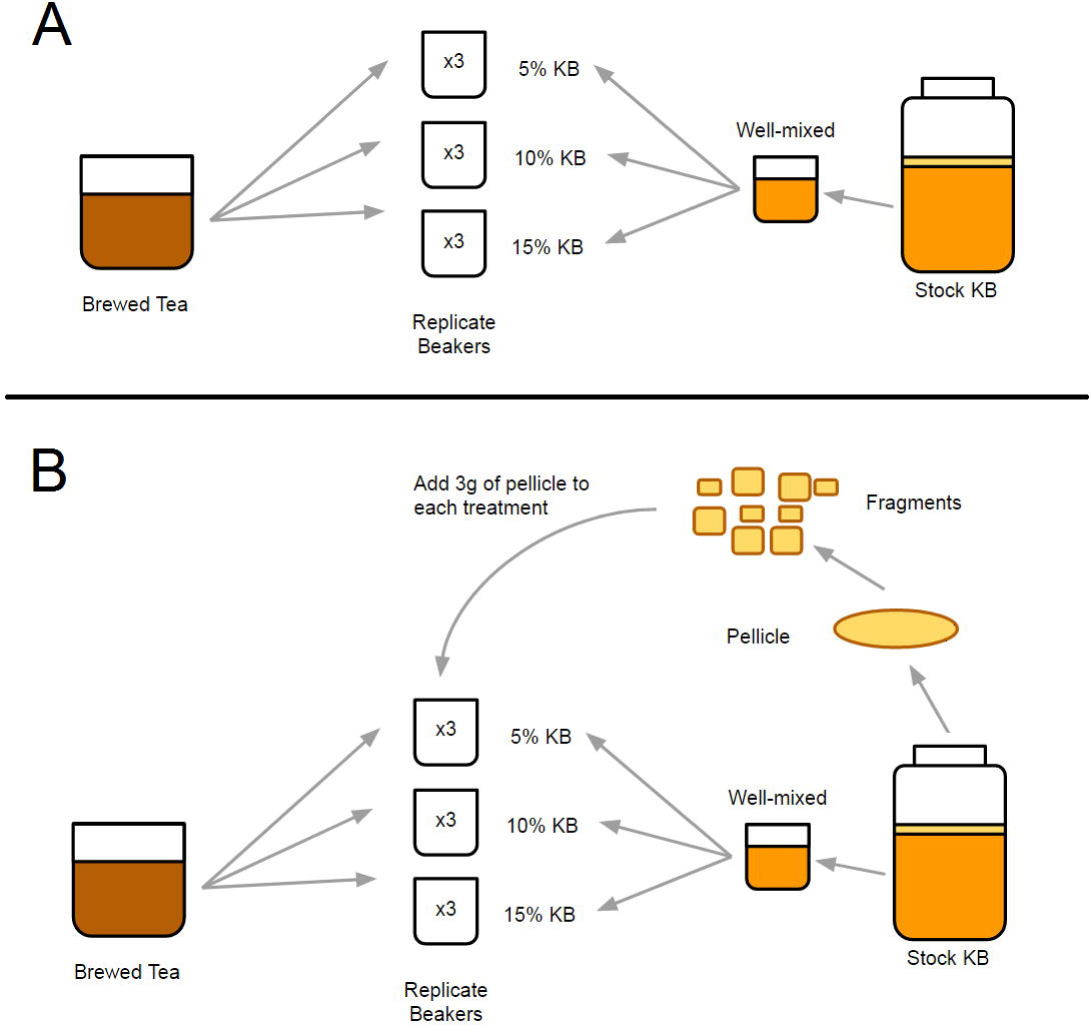
(made with Google Slides). Schematic of the experimental design for Experiment 1, showing the 2 arms of treatments: Liquid-only (A) and Liquid-and-pellicle (B) (see Methods).

Additionally, another set of 3 replicates were prepared that contained these same proportions of liquid (5% stock mix, 10% stock mix, and 15% stock mix) but also included fragments of pellicle from the stock vessel totaling 3g. These fragments were prepared by aseptically removing the topmost pellicle from the stock vessel (six weeks from last STM replenishment) by hand using 70% ethanol-sterilized gloves and placing it onto a plastic cutting board that had been sterilized with a 10% (v/v) bleach-DI water solution (after the board had dried). The pellicle was thoroughly rinsed 3 times with DI water, and then dabbed with a paper towel on both sides. Average pellicle thickness was measured with digital calipers (Fowler Company Inc., MA) at 4 equidistant points 1cm from the edge of the pellicle in addition to 1 point at the center. The average thickness of the pellicle fragment added to each replicate was 3.31mm (SD: 0.51mm). 2cm by 2cm square fragments were cut from the pellicle using an autoclave-sterilized straight razor and pooled together. Each beaker in these treatments received a random assortment of these fragments weighing 3g in total. All 6 treatments consisting of 3 replicates each were then placed into a 30°C incubator (Model 12E, Quincy Lab Corporation, IL) for 18 days, at which point they were harvested.

For the harvest, the entire mass of pellicle was removed from each replicate beaker using tweezers sterilized by a 70% ethanol solution. They were rinsed with DI water and dried as described in the experiments above, placed onto a large polystyrene weighing boat (Cole-Parmer, IL), and then weighed on an Accuris Instruments precision balance (Chemglass Life Science, NJ) to obtain a measure of fresh biomass weight. To keep a representative sample for future studies, a small fragment of each pellicle was removed with a razor, weighed separately, and then placed into a -20°C freezer. The remaining large pellicle fragments were then placed into a 30°C incubator and allowed to dry for 24 hours, at which point they were weighed again. The ratio of wet to dry weight for each large pellicle fragment was used to calculate the dry weight of the associated small frozen fragment. Both the recorded large fragment weight and calculated small fragment weight were summed to produce a total dry weight measure for each pellicle.

### Pellicle Consistency Experiment

The goal of this experiment was to determine whether homogenizing the pellicle could lead to more consistent results in the generation of new kombucha cultures (Fig 6). 3 treatments of 4 replicate beakers were prepared, all containing the same proportion of stock solution to STM (10%, as prepared in experiment 1, totalling 400mL), with 1 treatment as a liquid-only condition, 1 treatment supplemented with 0.5g of fragments of pellicle and the third treatment supplemented with 0.5g of homogenized pellicle. In order to standardize the extraction of pellicle fragments from the stock pellicle, the pellicle was divided into a grid of 2cm by 2cm squares. After washing and drying as in experiment 1, a randomly-generated number was used to select squares to extract fragments using a 10mm biopsy punch (AcuPunch, Ft. Lauderdale, FL). These pellicle fragments were collected until their total fresh weight was 0.5g, at which point they were added to the replicate beakers in this treatment. The final treatment consisted of collecting 2g of biopsy punched fragments from randomly assigned squares of the same grid-marked pellicle, as in second treatment. These fragments were added to a NJ600 Blender (SharkNinja Operating LLC, Champlain, NY) along with 1600mL of 10% stock-STM solution, and blended at ‘1’ power for 10 seconds and then ‘3’ power for 10 seconds. 400mL of the pellicle slurry was then distributed equally to 4 replicate beakers. Incubation and harvesting proceeded as in experiment 1.

**Figure 6.**
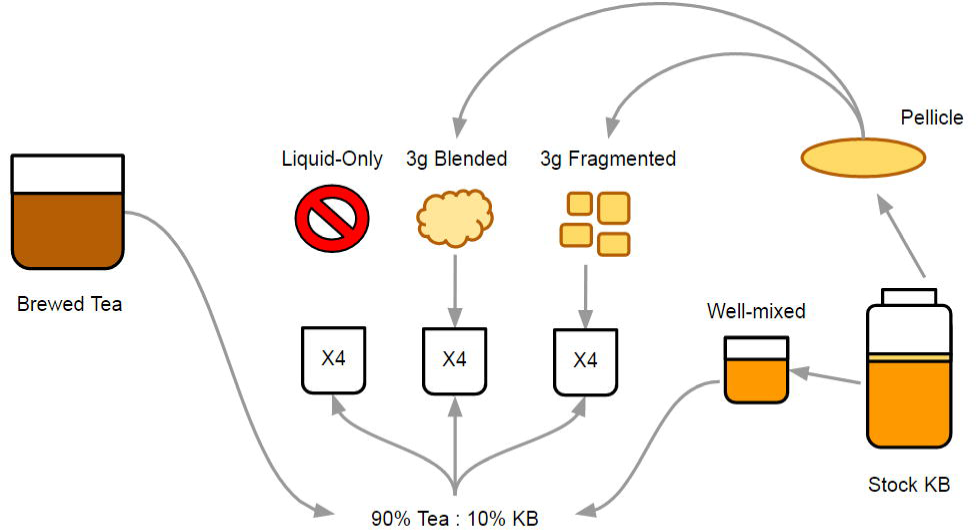
(made with Google Slides). Schematic of the experimental design for Experiment 2 (see Methods).

### Data Analysis

R statistical software (version 3.1, 2014) was used for all data analysis. We explored the data graphically using program-generated boxplots and scatterplots, and proceeded to examine linear correlations using the Spearman rank-order correlation coefficient. Wet and dry pellicle weights were highly correlated (ρ=0.9071), thus dry weight was used as primary dependent variable for analysis. Levene’s test was performed to determine the nature of variance between treatments, and data were natural-log transformed to ensure homoscedasticity. For experiment 1, treatments were analyzed using a paired-samples t-test using dry weight as the variable of interest. A 1-way ANOVA was then performed on these data with the presence of pellicle as the factor. For experiment 2, variance was analyzed using Levene’s test and the data was natural-log transformed as in experiment 1. A 1-way ANOVA was performed on these data with treatment as the factor.

## Acknowledgments

The authors wish to thank Arvind Varsani, Helen Wasielewski, Pamela Winfrey, Chaya Fux, Diego Mallo Adán, the Microbiome and Behavior Project members, and the Cooperation and Conflict lab for their valuable input and insights during the development of the project.

This research was supported in part by National Institutes of Health grants P01 CA91955, R01 CA149566, R01 CA170595, R01 CA185138 and R01 CA140657 as well as Congressional Directed Breast Cancer Research Program Award BC132057, and by a grant from the John Templeton Foundation titled “Generous by nature: Need-based transfers and the origins of human cooperation” (to A.A.). The findings, opinions and recommendations expressed here are those of the authors and not necessarily those of the universities where the research was performed, the John Templeton Foundation, or the National Institutes of Health.

The authors declare that they have no conflicts of interest with any products or equipment used during the course of this project.

